# Social-ecological models with social hierarchy and spatial structure applied to small scale fisheries

**DOI:** 10.1101/2024.02.17.580847

**Authors:** Sophie Wulfing, Easton R. White

## Abstract

Socio-ecological models combine ecological systems with human social dynamics in order to better understand human interactions with the environment. To model human behavior, replicator dynamics can be used to model how societal influence and financial costs can change opinions about resource extraction. Previous research on replicator dynamics has shown how evolving opinions on conservation can change how humans interact with their environment and therefore change population dynamics of the harvested species. However, social-ecological models often assume that human societies are homogeneous with no social structure. Building on previous work on social-ecological models, we develop a two-patch socio-ecological model with social hierarchy in order to study the interactions between spatial dynamics an social inequity. We found that fish movement between patches is a major driver of model dynamics, especially when the two patches exhibit different social equality and fishing practices. Further, we found that the societal influence between groups of harvesters was essential to ensuring stable fishery dynamics. Next, we developed a case-study of a co-managed fishery where one group fishes sustainably while another was over-harvests, resulting in a fishery collapse of both patches. We also found that because social influence only included number of fishers and not effective strategies, increased social pressure actually decreased the sustainability of the fishery. The findings of this study indicate the importance of including spatial components to socio-ecological models and highlights the importance of understanding species movements when making conservation decisions. Further, we demonstrate how incorporating fishing methods from outside sources can result in higher stability of the harvested population, indicating a need for diversified information when managing resources.

## 2 INTRODUCTION

Social ecological models treat human behavior as a variable as opposed to a set parameter. Allowing human heavier to be dynamic allows for the study of how human decision making can change in response to environmental factors and, in turn, alter how humans interact with resources and profits (Bauch, 2005; Ostrom, 2009; Innes et al., 2013; Oraby et al., 2014; Bauch et al., 2016; Sigdel et al., 2017; Thampi et al., 2018). As human societies grow increasingly intricate and interconnected, these models can help us to analyze how our social structures can influence the environment around us (Liu et al., 2007). Social ecological modeling provides important insight not only into how human decision making can influence ecological patterns but can also show hidden processes, reveal regime shifts that would otherwise be hidden, and identify vulnerabilities of systems that do not exist within the purely social or ecological models (Liu et al., 2007; Young et al., 2007; Ostrom, 2009; Lade et al., 2013). Socio-ecological models can also be used in systems where data are difficult to collect, as parameters can be changed in order to analyze different hypothetical scenarios. Conservation plans often do not reach their conservation goals, and these setbacks are often attributed to a lack of stakeholder participation (Crona & Bodin, 2006; Salas et al., 2019; Prince et al., 2021). Socio-ecological models can identify where areas of potential conflict can arise, compromises that can be made in the system, and alternative conservation practices that encourages participation from all stakeholder groups (Ban et al., 2013). Further, as social-ecological models are simulations of human and environmental interactions, they allow flexibility and can be adapted to fit the specific system of study and improve place-based management practices (Young et al., 2007; Liu et al., 2007; Felipe-Lucia et al., 2022)

Due to their adaptability, socio-ecological models can use a wide range of strategies to represent human decision making. One such method is replicator dynamics, which model human decision making where an individual makes conservation choices based on weighing the perceived benefits of conservation with the costs, as well as the social pressure to conform to the group’s stance on conservation (Bauch, 2005). Individuals will therefore “replicate” the behavior of their peers by changing their harvest practices based on the opinion of the majority (Bauch & Bhattacharyya, 2012). Models that employ replicator dynamics have been used to show how this social learning is a key component to vaccination uptake in public health, and preexisting social norms can actually suppress vaccine uptake despite frequent disease outbreaks (Bauch & Bhattacharyya, 2012; Oraby et al., 2014). Replicator dynamics can also have conservation applications as pest invasion models have shown ways to simultaneously mitigate pest outbreaks and the cost to address them in the timber industry (Barlow et al., 2014). Further, land use changes have been modeled to have completely different dynamics when human decision making was added to replicator dynamic models (Innes et al., 2013). However, past work on human behavior has generally assumed that human societies are homogeneous, and all people are subject to the same social influence and ecological dynamics.

Instituting effective conservation strategies can be especially difficult if the organism being protected has a migratory pattern that crosses over multiple management jurisdictions such as country borders (Ogburn et al., 2017; Garrone-Neto et al., 2018; Ramírez-Valdez et al., 2021). Borders can also create challenges when gathering population data that require extensive fieldwork (Cozzi et al., 2020; Hebblewhite & Whittington, 2020). The fragmentation of management can also result in a mismatch of conservation strategies that become ineffective when the distinct management bodies do not coordinate efforts (Siddons et al., 2017). Research on the importance of coordinated research efforts has been conducted on many terrestrial species with large migratory ranges and have consistently shown that cooperation among government bodies is essential to protecting the health of highly migratory species or species whose native ranges expand across multiple countries (Plumptre et al., 2007; Gervasi et al., 2015; Meisingset et al., 2018). Because fish are generally migratory, management coorporation is especially relevant in international waters or waters where different government bodies share jurisdiction (Mchich et al., 2000). Previous research on two-patch fishing models has shown that movement rates between patches can affect population stability when there are different fishing pressures in each patch (Mchich et al., 2000; Cai et al., 2008). Economic output can also be maximized in multi-patch fishing models as high dispersal can result in a higher overall yield of the system than the yield of each patch combined (Auger et al., 2022). High dispersal across patches is commonly found to be an essential component to maximizing population health and economic gain from fishing (Freedman & Waltman, 1977; Moeller & Neubert, 2015; Auger et al., 2022). Two-patch models help us to understand the population dynamics of fish species better who face different pressures in each patch and have even resolved conflicts between fishing groups (Mchich et al., 2000).

Contrary to the assumption made by previous models that human groups are homogeneous, the vast majority of real-world societies exhibit some form of hierarchy or inequality. Societies with different social subgroups can often exhibit an “us vs. them” mentality and compete for resources (Borgatti, 2003). People’s relationship with the environment has been shown to be influenced by many factors such as social status, wealth, gender, education, and even notions of self-importance (Baker-Médard, Gantt, et al., 2021). Competition over resources has been shown to be exacerbated by social hierarchies and ‘top-down’ regulation whereas when social connectivity is considered in management plans, management outcomes are not only improved, but costs are reduced as well (Krackhardt & Stern, 1988; Grafton, 2005; Bodin & Crona, 2009). Further, members of social networks have been shown to have varying levels of connectivity with others in their groups based on attributes such as ethnicity, which can in turn alter an individual’s relationship with the environment and their views on conservation (Barnes-Mauthe, 2013; Sari et al., 2021). Barnes-Mauthe (2013) showed that fishing communities can exhibit homophily, which is the tendency for people to obtain information and opinions from those who are similar to themselves before seeking views from those who are perceived as different. Therefore, people in different social groups may be receiving different information and opinions about conservation and acting accordingly (McPherson et al., 2001). For example, in Kenya, communication among fishers has been shown to stay within groups using the same gear type which has inhibited successful regulation of the whole fishery (Crona & Bodin, 2006). Further, in the southwest Madagascar octopus fishery, fishing method and location typically falls along gendered lines. When fishing restrictions were imposed on tidal flats, women’s access to octopus harvest was restricted, while men, who typically fished in deeper waters, were able to maintain their livelihood (Baker-Médard, 2017). In Thailand, ethnicity has been shown to be a source of fishing conflict which has exacerbated resource depletion (Pomeroy et al., 2007). The existence of social structures is extremely prevalent in human societies which can affect how people interact with the environment. However, there is little existing research that uses replicator dynamics study to study how social hierarchies alter harvest practices.

Small-scale fisheries are a particularly relevant system to apply replicator dynamics as fishing practices and policies are often made by communal decision makers. Research on small-scale fisheries is a growing and essential field as they are drastically understudied yet affect many people around the globe (The World Bank, 2012). Due to tight social structures, community decision making and strong reliance on the environment, small-scale fisheries are systems that are well represented by socio-ecological models and replicator dynamics (Grafton, 2005; Thampi et al., 2018; Barnes et al., 2019). Governmental bodies or third parties instituting conservation efforts in small-scale fisheries have often been unsuccessful, especially when the social and economic components of the industry have been ignored (Salas et al., 2019; Prince et al., 2021). However, even when human interactions and decision making have been considered, socio-ecological models have often treated individuals in human societies equal in their social standing. As human societies are often complex and hierarchical, the simplistic assumption that everyone interacts with the environment and within their community equally can lead to lack of participation in conservation by some groups within a community (Barnes-Mauthe, 2013; Cumming et al., 2017). Mismanagement of fisheries have even been shown to exacerbate these social inequalities (Cinner et al., 2012; Baker-Médard, 2017). Further, the specific dynamics of the fishery in question have been shown to be important components to models, as models with multiple patches can actually mitigate over-fishing if there is high movement of the harvested species between patches (Cressman et al., 2004). No previous research has combined two-patch fishing models with a hierarchical human decision making model in order to study how space and social dynamics affect fishery dynamics.

In this study, we couple a human-decision replicator dynamics model with social hierarchies with a two-patch resource model in order to understand how decision making is affected by spatial and hierarchical factors. The objectives of this study were: 1) to compare the output of previous replicator dynamics studies with the new two-patch model to understand the affect of species movement on harvesting decisions, 2) understand the effect of social hierarchy and communication across groups on the dynamics of this model, 3) use a co-managed small-scale fishery as a case study to understand how fishery dynamics are driven when one group fishes sustainably while the other over-harvests. We hypothesized that higher cooperation between groups would benefit fish stocks overall and that increased fish movement would increase the health of fish populations.

## 3 METHODS

### 3.1 Model Construction

We build on the work of Bauch et al. (2016) by extending their old-growth forest model to a two-patch model (Figure 1). The resource population models adapted from Bauch et al. (2016) are as follows:

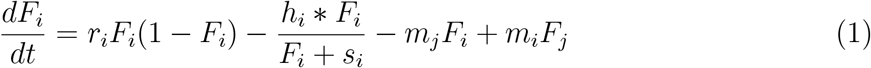

where the change in resource populations *F*_*i*_ is dependent on *r*_*i*_, the net population growth of each patch *i*, and both populations follow logistic growth. The second term: 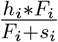, denotes population lost to human activity. *h*_*i*_ is the harvesting efficiency of the respective human population and *s*_*i*_ controls the supply and demand of the system. Because we extend this to a two-patch model, the *m*_*i*_ parameter denotes the movement of the harvested species out of patch *i* and into patch *j*. In this study, we assume a closed population between the two patches. Therefore, individuals move directly from patch to patch and do not disperse elsewhere, nor are individuals immigrating from outside areas.

**Figure 1:**
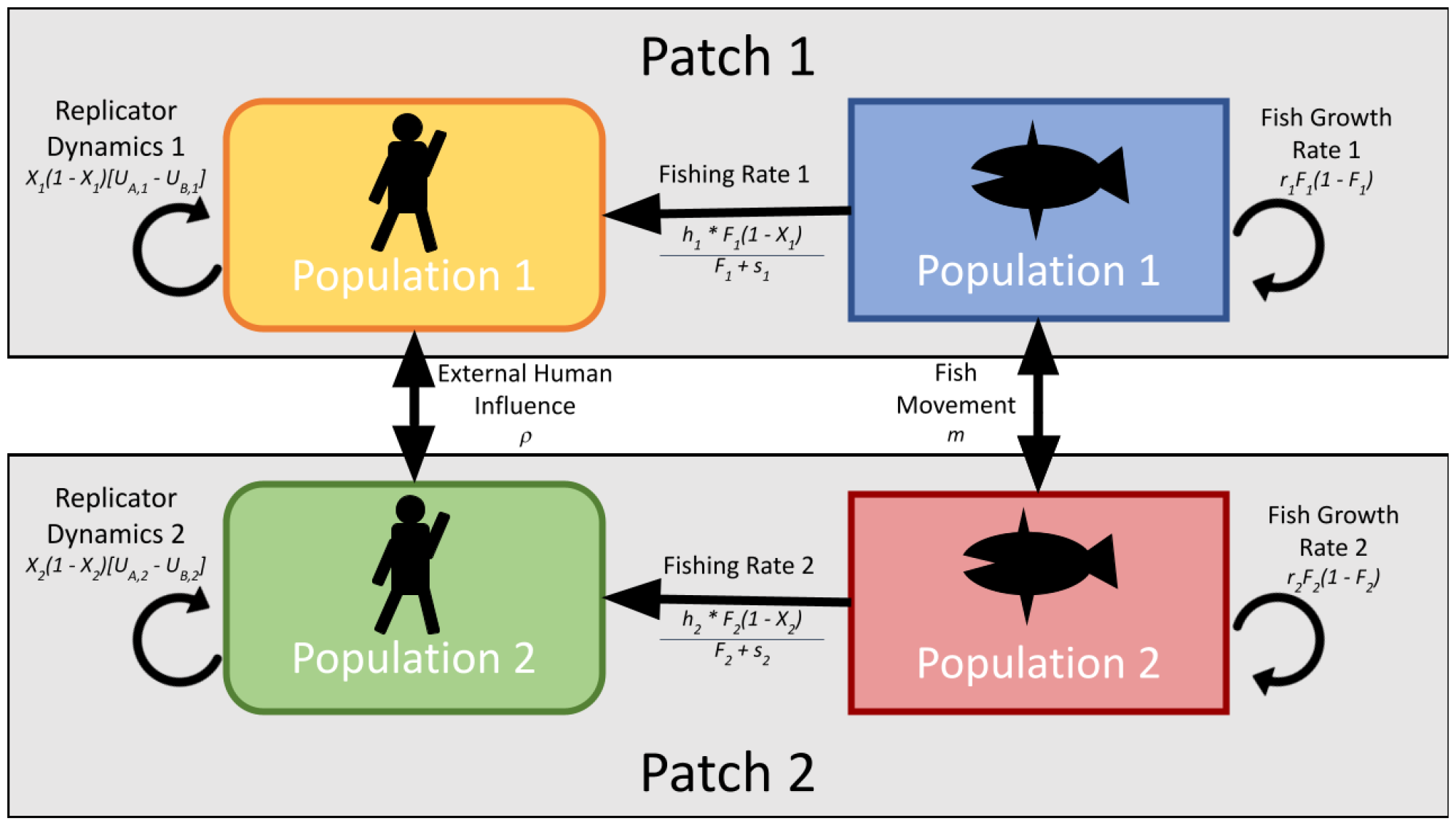
A conceptual representation of our model as a two-patch extension of Bauch et al. (2016). Here, each fish population (*F*_*i*_) in each patch *i* increase through natural growth and movement of fish into the patch. Fish populations are decreased through emigration out of the patch and fishing mortality. The number of fishers (*X*_*i*_) in each patch *i* change in response to fish population levels, the cost of stopping fishing activity, and the opinions of those in the patch and those in the other patch.

For the model of human activity and opinion, we use replicator dynamics from evolutionary game theory to simulate societal influence on an individual’s opinion. Humans in this population can either be harvesters (therefore participating in harvesting activity) or conservationists (who do not partake in resource extraction), but can change from their current opinion to the other based on the perceived values and costs of each stance. Social dynamics are represented by the proportion of conservationists in a population (*X*) and the proportion of harvesters (1 − *X*). These two groups of conservationists and harvesters interact with one another using the term (*X*)(1 − *X*) which simulates individuals “sampling” the opinions other individuals in the population. If one opinion dominates in the population (i.e. *X >>* (1 − *X*) or (1 − *X*) *>> X*), the rate of changing opinions will be slow as the power of societal pressure makes it challenging for the other opinion to gain traction. However, if *X* and (1 − *X*) are close, the rate of change in opinion will be fast as society has a split opinion on conservation versus harvest, so individuals will be quick to take up the opinions of others. In this model, each person holds an opinion (conservation or harvest) by weighing the benefits of conservation (*U*_*A*_) against the benefits of harvest (*U*_*B*_), resulting in the replicator equation:

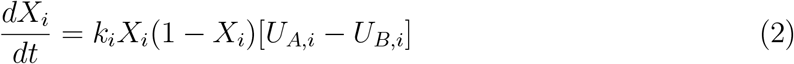

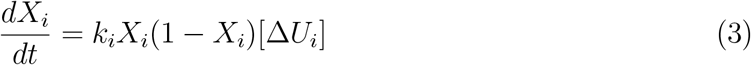

where *k*_*i*_ refers to the rate of interaction within a group. As individuals “sample” the opinions of others in their group, they can switch from A to B if *U*_*B*_ > *U*_*A*_ and vice versa. In our model, we adapted *U*_*A*_, the perceived benefit of conservation, from Bauch et al. (2016) with the added influence of the other population’s opinion. *U*_*A*_ is therefore given by:

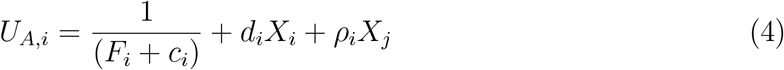

where 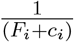 represents the perceived rarity of the harvested population within a patch. As *F*_*i*_ and *c*_*i*_ (the rarity valuation parameter) decrease, perceived rarity will increase, therefore adding to the perceived benefit of protecting resources. *d*_*i*_ refers to the social influence that each population has on itself, and as an individual encounters a conservationist in their own population (*X*_*i*_), the social benefit of also being a conservationist is shown in *d*_*i*_. *ρ*_*i*_ has this similar effect of social influence, but denotes the social effect of the opposite population on decision making (*X*_*j*_). Individuals in each population *i* are receiving information about the conservation practices of the other population *j*, and the influence that this has on each population is encapsulated by *ρ*_*i*_.

*U*_*B*_ (the perceived benefits of harvest) is:

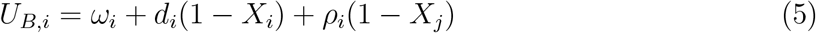

where *ω*_*i*_ is the cost of conservation (i.e. revenue lost by not harvesting) where now, *d*_*i*_ is the within-population social benefit of switching to harvesting (1 *− X*_*i*_) and *ρ*_*i*_ is the other population’s (1 *− X*_*j*_) ability to change the opinion of an individual to be a harvester.

Plugging equations (4) and (5) into equation (2) gives:

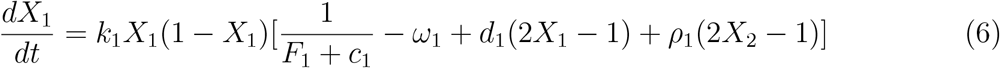

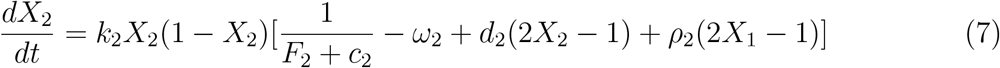

where specifics of the derivation are outlined in the supplementary material. Coupling the resource population and human opinion models gives:

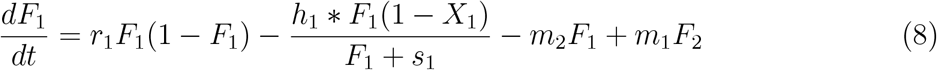

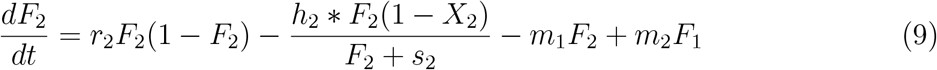

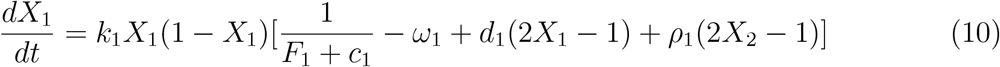

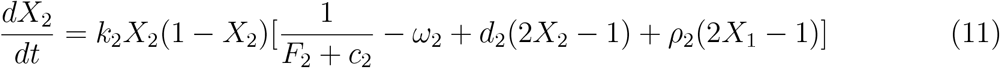

where the harvesting pressure is now a function of the number of harvesters in a population 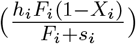. Further, the opinion of each population will shift based on the perceived population health of their respective patch weighed against the costs and benefits of conservation. As resources decrease, individuals will sway more toward conservation, thereby relieving harvest pressure. However, we now have an external influence in this model: the opinions of people in population *j*. The strength of this external influence is *ρ*, and in this study, we plan to simulate inequalities in human societies with this parameter.

The default parameters used to analyze the resources movement and human hierarchy parameters were taken from an analyses done in Bauch et al. (2016) and given in Table 1. Here, Bauch et al. (2016) found an oscillatory behavior where decreased forest cover resulted in decreased harvest due to the replicator dynamics of the human system which allowed for forest recovery and humans to begin high harvest once again.

**Table 1:**
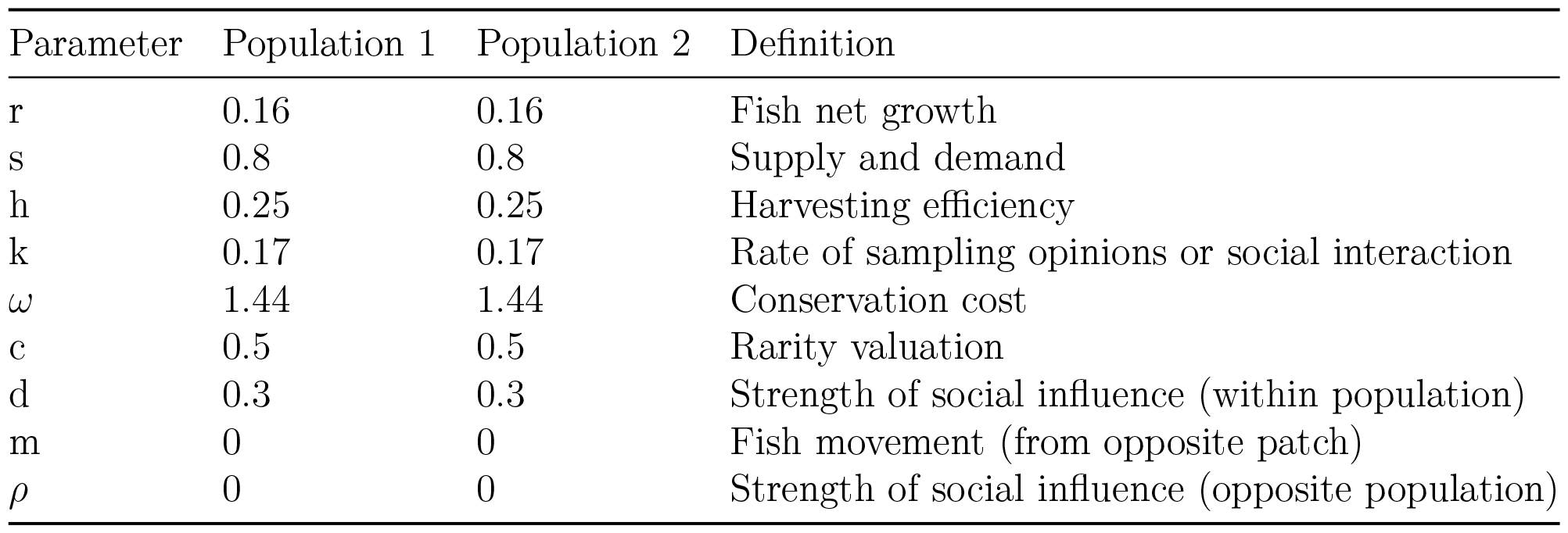
Default parameter values used in this analysis taken from Bauch et al. (2016) where oscillations are observed.

### 3.2 Parameter Analyses

In order to understand how resource movement (*m*_1_ and *m*_2_) affects dynamics, we first compare how the system will change when both patches are equal (i.e. all of the parameters in each patch is the same) by increasing both *m*_1_ and *m*_2_ incrementally and running the model for 1000 years. We then compare this to the asymmetrical case, where we just increase the *m*_1_ parameter and see the effect on the model for the next 1000 years. We also construct bifurcation curves of the *m*_1_ parameter when compared to resource populations in order to understand their effect on dynamics. Further, to analyze the human hierarchy parameters *ρ*_1_ and *ρ*_2_, we constructed this same analyses of increasing *ρ*_2_, or the amount of influence of human population 2 (*X*_2_) has on the dynamics of human population 1 (*X*_1_). We also compared this to the effect on incrementally increasing *d*_1_.

### 3.3 Co-Managed Small Scale Fishery Case Study

For a small scale fishery, we choose to model a two-patch fishery where patch 1 is fishing sustainably while patch 2 is over-harvesting. The harvested fish species has a mid-range growth rate and regularly diffuses across the two patches, such as the parrot fish modeled in Thampi et al. (2018), which uses a fish growth rate of is 0.35 fish per year, but alter patch 1’s growth rate to be 0.4 fish per year. For the harvesting efficiency, we choose a maximal fishing rate of 0.5. These parameters were adapted from a coral reef fishing model Thampi et al. (2018) where *r* = 0.35 and *h* = 0.5 are the mid-level growth rate and max fishing rates analyzed by this paper. For the movement parameters *m*, we chose 0.2 for each as these are the values used in the two-patch fishing model described in Cai et al. (2008). We used the *s* parameter described in the Bauch et al. (2016) model of *s* = 0.8. For the purposes of our study, we are assuming a constant net growth rate of fish populations and that reproduction happens locally within each patch. The rate at which humans interact with one another is described by the parameter *k*. In our default model, we use *k* = 1.014 as adapted from the Thampi et al. (2018) default model. Thampi et al. (2018) calculated this parameter by fitting conservation opinion data in the United States from 1965 to 1990 to coral health data at that time (Thampi et al., 2018). We used the default rarity valuation parameter *c* from Thampi et al. (2018) where *c* = 1.68. The cost of conservation default parameter is *ω* = 0.35 from Bauch et al. (2016). Further, as our default model has no human social hierarchy, we set *d* = *ρ* = 0.5 for our social norm strengths as adapted from Bauch et al. (2016) which models social decision making regarding deforestation.

Based off of the default model described above, we then change parameters such that patch 1 is fished sustainably, meaning the fish population in patch 1 is able to persist regardless of the fishing pressure from human population 1. We then set patch 2 to be over-fished, meaning human patch 2 is fishing at too high a rate for the fish population to survive over time (Table 2). Further, we add a social hierarchical component where patch 2 has a higher social influence on patch 1. To analyze the overfishing scenario, we incrementally increase the parameters *m* and *ρ* and simulated this system for 100 years in order to assess how increasing each new parameter would affect the overall dynamics of the system.

**Table 2:**
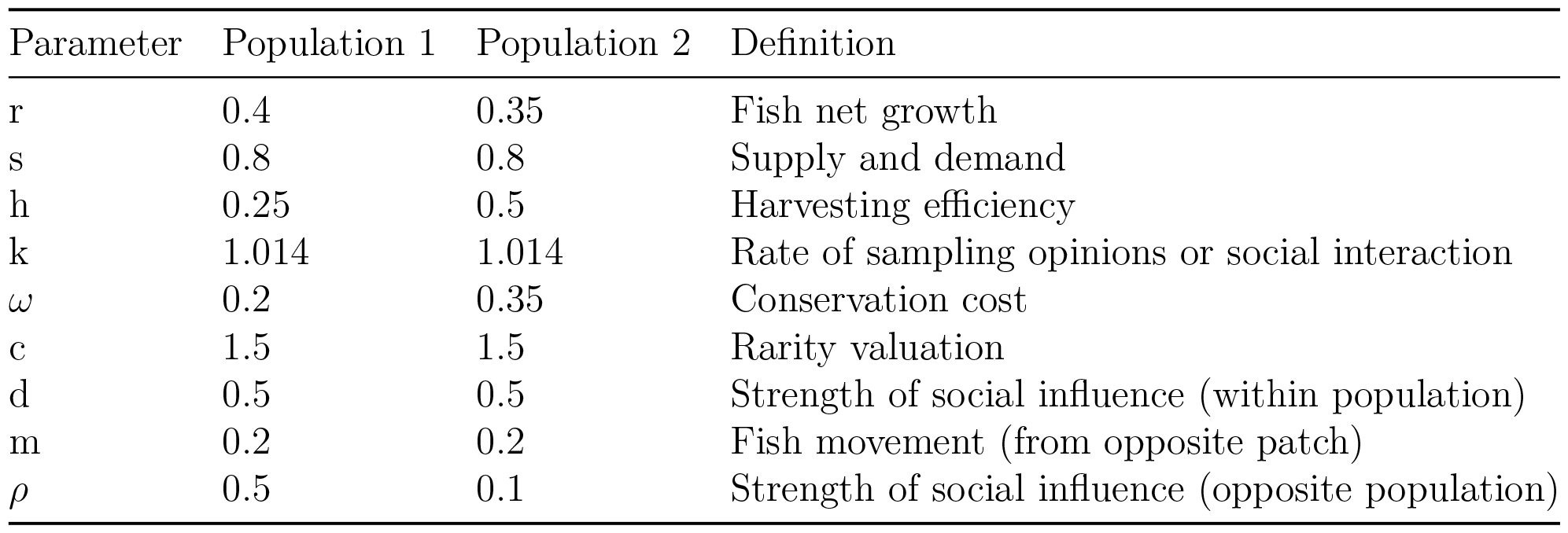
Parameter values used to simulate sustainable fishing practices in patch 1 and over-fishing in patch 2.

| Parameter | Population 1 | Population 2 | Definition |
| --- | --- | --- | --- |
| $r$ | 0.4 | 0.35 | Fish net growth |
| $s$ | 0.8 | 0.8 | Supply and demand |
| $h$ | 0.25 | 0.5 | Harvesting efficiency |
| $k$ | 1.014 | 1.014 | Rate of sampling opinions or social interaction |
| $\omega$ | 0.2 | 0.35 | Conservation cost |
| $c$ | 1.5 | 1.5 | Rarity valuation |
| $d$ | 0.5 | 0.5 | Strength of social influence (within population) |
| $m$ | 0.2 | 0.2 | Fish movement (from opposite patch) |
| $\rho$ | 0.5 | 0.1 | Strength of social influence (opposite population) |

## 4 RESULTS

### 4.1 Movement Parameter

To analyze the result of space on socio-ecological models, we observed the effects of increasing both *m*_1_ and *m*_2_ simultaneously (the symmetrical case) and compared this to the effects of only increasing *m*_1_, or the movement of resources from patch 2 to patch 1 (Figure 2). Here, we find that movement does not change dynamics in the symmetrical case (Figure 2 a), b), and c)), showing that if all parameters are the same in each patch, the movement of resources between them does not change dynamics. However, if there are differences between patches (Figure 2 d), e), and f)), resource movement will greatly alter dynamics and if the model is undergoing oscillations, the linear aspects of the movement parameters will eventually overcome the non-linear dynamics of oscillations if the movement parameter is sufficiently high.

**Figure 2:**
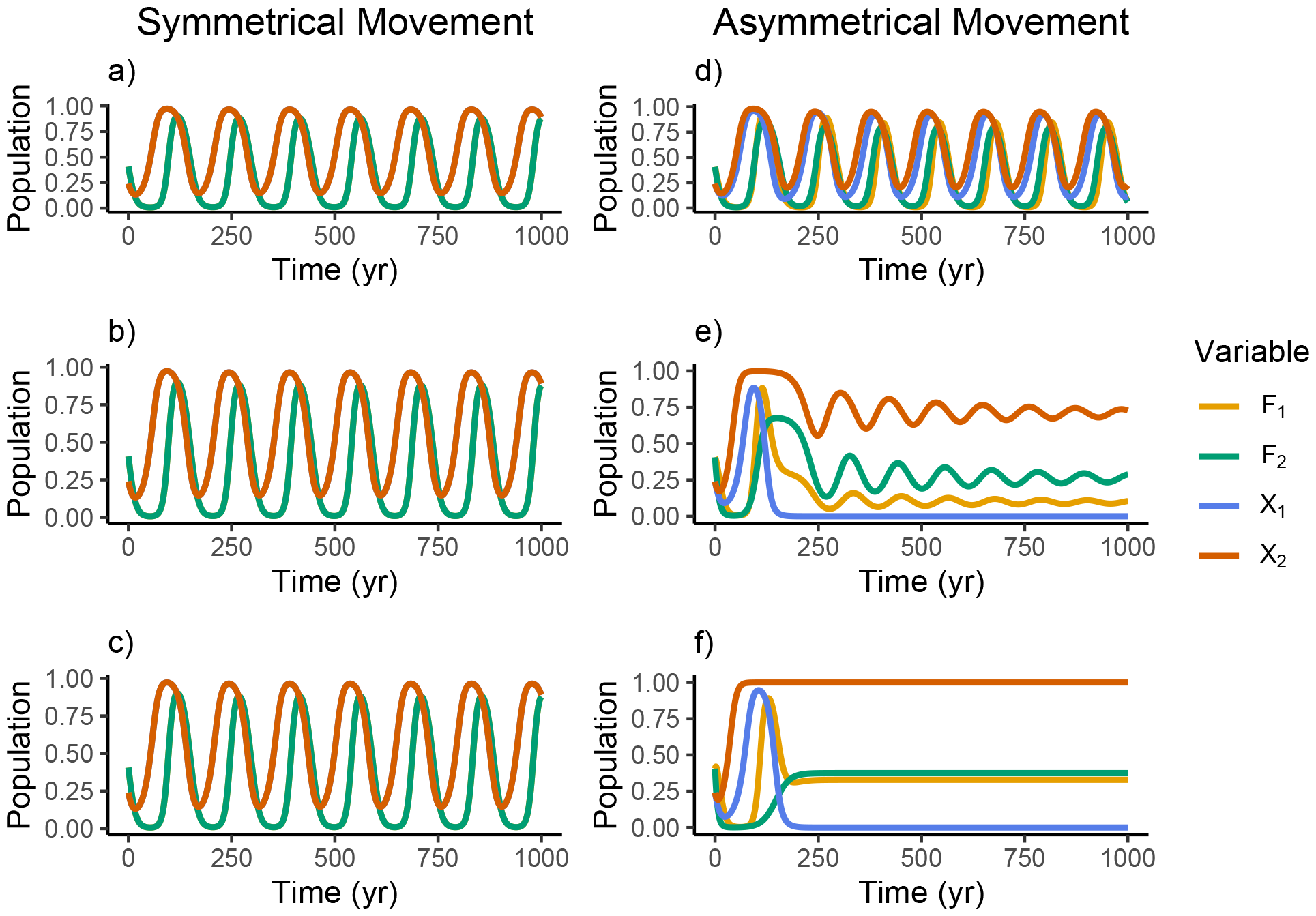
In graphs a), b), and c), both *m*_1_ and *m*_2_ were set to 0.01, 0.05, and 0.1, respectively. The corresponding graphs show the dynamics of these models with the new parameterizations. d), e), and f) show the changes in model dynamics when *m*_2_ is held at 0 and only *m*_1_ (the movement of resources from patch 2 to patch 1) is increased by 0.01, 0.05, and 0.1, respectively. All other parameters were held at the values given in table 2

### 4.2 Social Hierarchy Parameter

In figure 3, we can see that increases in *d*_1_ result in higher amplitude oscillations, where *F*_1_ will dip to almost 0 for many years then recover back to 1. Increases in *d*_1_ affect the model differently than increases in *ρ*_2_, the influence of the other human population. Here, the population dynamics of *F*_1_ stay relatively constant around 0.2, and only have very small oscillations around this number, therefore increases in *d*_1_ can result extreme booms and busts of resource populations while increases in *ρ*_2_ results in limited populations, but these but the resulting dynamics oscillate less, which indicates more stable dynamics. Increases in either *d*_1_ or *ρ*_2_ result in less frequent oscillations, meaning humans are slower to change population levels and that plot 1’s resource populations spend more time at the peaks of their oscillations before either recovering from 0 or decreasing from 1.

**Figure 3:**
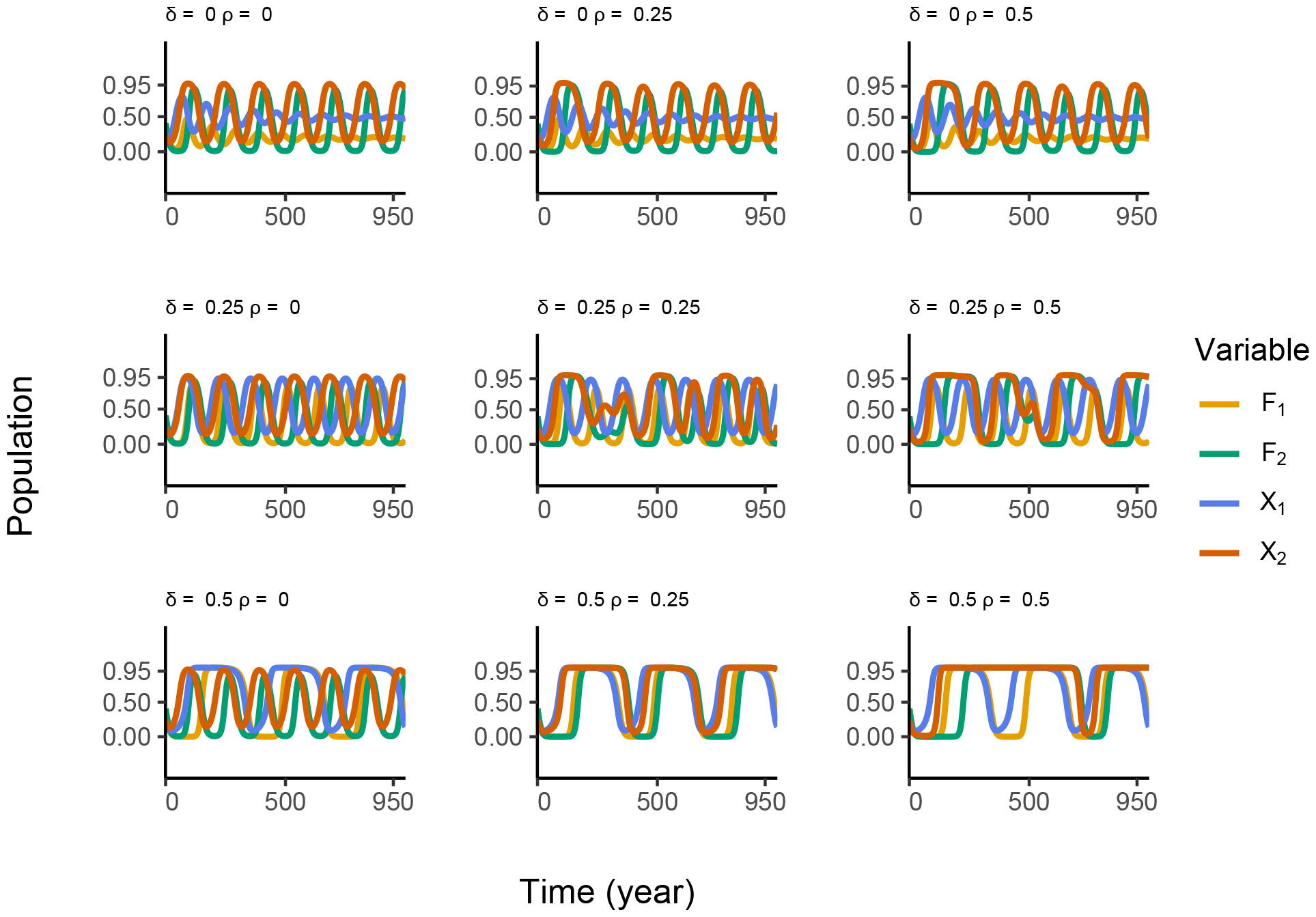
The difference in increasing social pressure within population 1 (the *d*_1_ parameter is increased down the columns of graphs) versus increasing social pressure from population 1 onto population 1 (the *ρ*_2_ parameter is increased across rows of graphs).

### 4.3 Scenario Analysis

We then modeled a hypothetical scenario where patch 1 is fished sustainably whereas patch 2 is experiencing over-fishing and has a higher social sway than patch 1. We modeled overfishing by altering fish new growth rates (r), harvesting efficiencies (h), costs of conservation (*ω*), and external social norm strengths (*ρ*) (Table 2). Here, the unsustainable practices of human population 2 are so exploitative, that both fish populations eventually collapse. We used this overfishing parameterization for the rest of the analysis of a co-managed small-scale fishery.

Next, we ran our model with the parameterization outlined in table 2 with incrementally higher external social influence values (*ρ*) in both populations and observed how this affected the final population of each fish patch (Figure 4). We found that under different parameterizations, there were often instances where *ρ* acted as a tipping point for population dynamics where instead of continuously changing the final fish populations, the *ρ* parameter either resulted in stable fish populations or both stocks collapsed once *ρ* increased past this tipping point.

**Figure 4:**
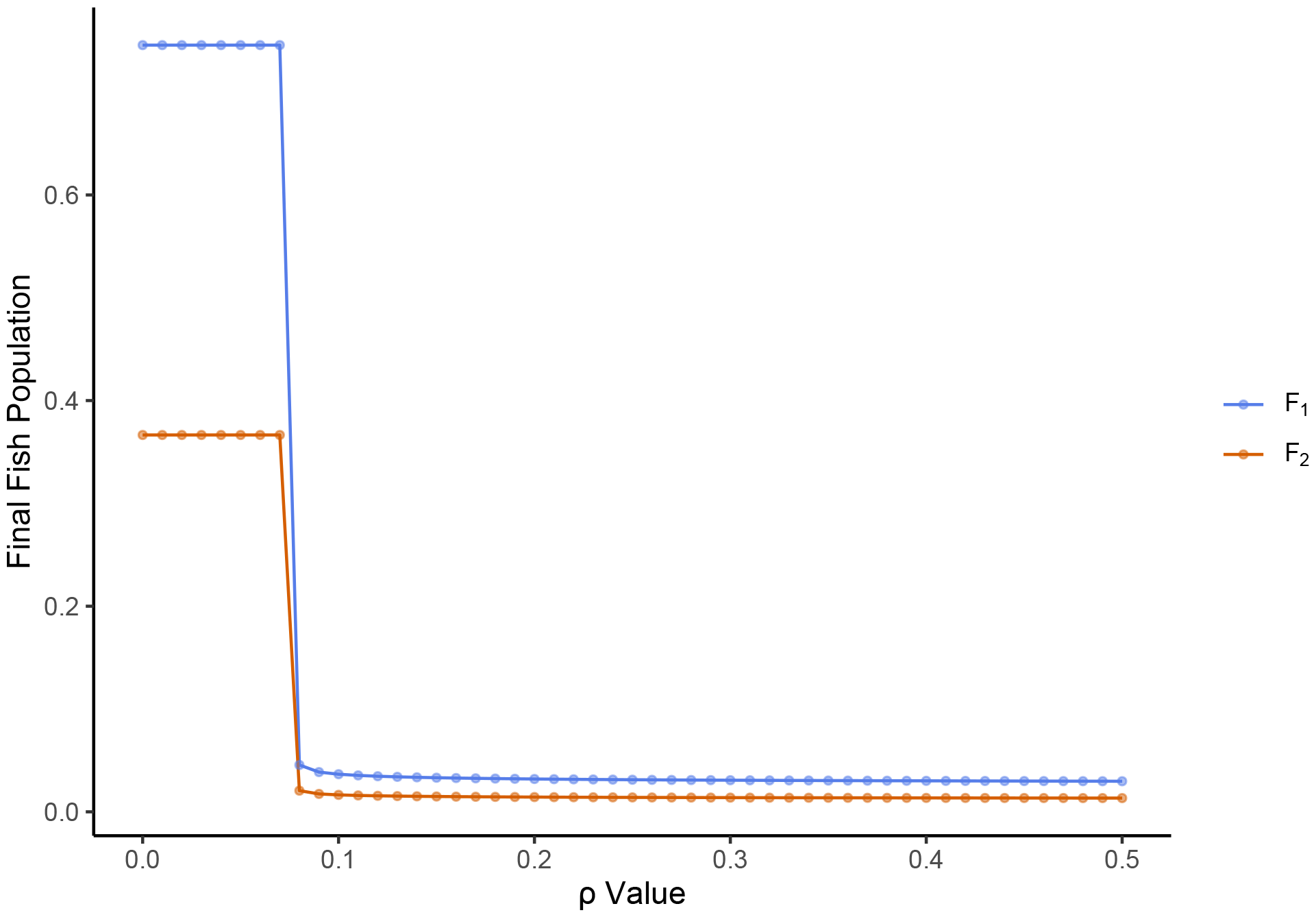
Final fish populations after 100 years in the two-patch fishing model where the *F*_1_ population in patch 1 is fished sustainably but human population 1 has a lower social influence than humans in patch 2, where *F*_2_ is being fished unsustainably. Both *ρ*_1_ and *ρ*_2_ were increased simultaneously.

We then ran the same analysis with the fish dispersal parameter, *m*, by changing *m*_1_ and *m*_2_ individually. Contrary to the effect external social influence (*ρ*) had on the model, dispersal had a more direct and continuous effect on the final population of fish in each patch. For example, as fish movement from patch 2 to patch 1 increased (i.e. from the unsustainable patch to the sustainable patch), this actually maintained low fish populations the sustainable patch, but resulted in crashed populations in the unsustainable (Figure 5 a). However, if the fish movement was increased from patch 1 to patch 2 (from the sustainable fishing to unsustainable), both patches eventually collapsed to zero (Figure 5 b).

**Figure 5:**
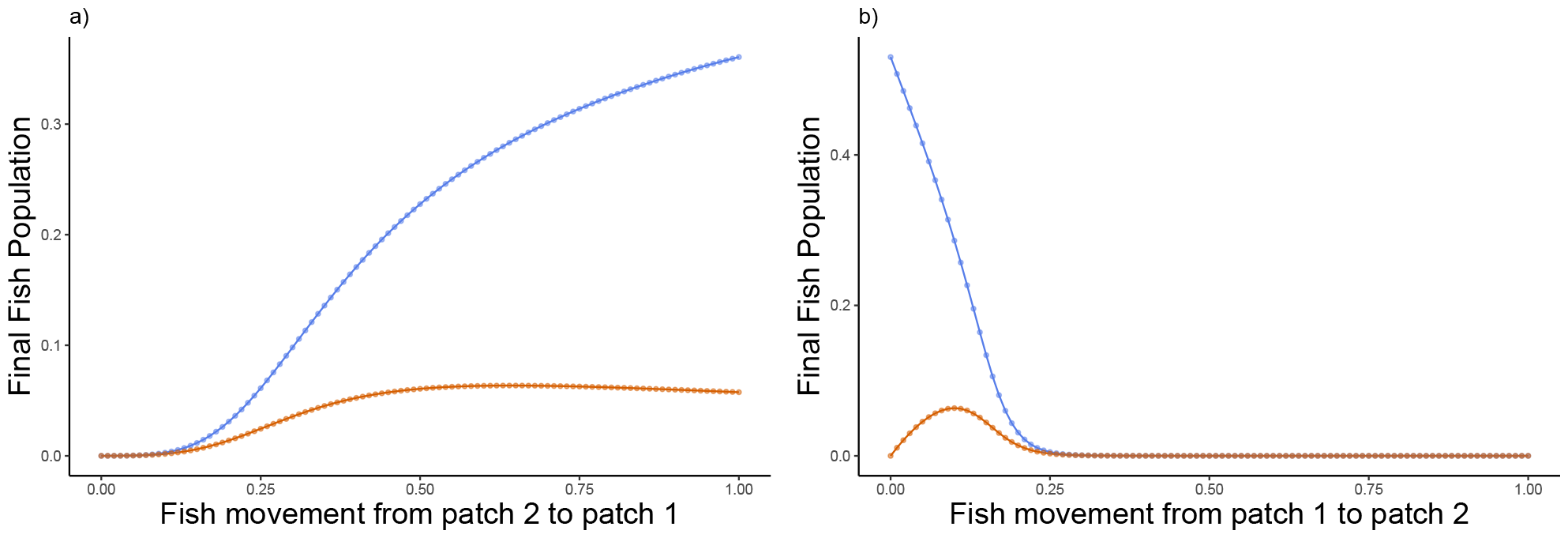
Final fish populations after 100 years in the two-patch fishing model where patch 1 (*F*_1_) is fished sustainably but human population 1 has a lower social influence than patch 2, where *F*_2_ is being fished unsustainably. a) shows how increases in fish movement into patch 1 (*m*_1_) affect final populations and b) shows how increases in fish movement into patch 2 (*m*_2_) affect final populations.

## 5 DISCUSSION

Instead of just social norms controlling the dynamics of our model, we found that the movement of the resource species between patches (m) was a major driver of population sustainability or collapse (Figure 5). As we increased the movement of fish into the sustainable patch in the fishery scenario (Figure 5 a), populations in that respective patch also increased because humans in population 1 continued to fish sustainably. Further, as those in population 2 decreased fishing rates, this influenced population 1 to also decrease their number of fishers. As a result, population 1 maintained high fish stocks while population 2 had low stocks. On the contrary, as fish moved from the sustainable patch 1 to the unsustainable patch 2 (Figure 5 b), both fish populations collapsed as *m*_2_ increased because fish movement away from patch 1 eventually grew to be too great for human population 1 to fish sustainably and human population 2 continued to over-fish in their own patch. When both patches are subject to the same conditions (Figure 2 a), b), and c)), resource movement does not affect the dynamics at all. It is only when each patch is subject to different conditions, in the case of figure 2 d), e), and f), where only the movement between patches is asymmetrical, does movement become extremely important in dynamics. This finding is especially relevant to co-managed fisheries, where different areas may be subject to different regulation, environmental conditions, or opinions about conservation. High migration has been shown to be an essential part of maximizing economic benefit from fishing in multi-patch models (Moeller & Neubert, 2015). Because fish are generally migratory and therefore can be difficult to track, constraining fishing to one group of people is more challenging (Grafton, 2005), especially for fish species that exhibit different movement patterns based on life stage, and requires more management coordination (Siddons et al., 2017).

The social hierarchy parameter *ρ* can also dictate whether or not patches will be harvested sustainably. Figure 3 exemplifies how, when increasing social susceptibility to one’s respective patch (increasing the *d* parameter), can result in booms and busts of resource availability. On the other hand, increasing social susceptibility to outside social influence (increasing the *ρ* parameter) can actually result in more stable dynamics because human population 1 is exhibiting a “portfolio effect” of harvest opinion. In other words, population 1 is taking in opinions regarding harvest from different sources, which can dampen extreme reactions to harvest decisions and therefore reduce extreme changes in fishing pressure. Portfolio effects have been shown to be beneficial when fishers diversify the species they catch, which allows them to compensate for lost catch when one species experiences decline (Finkbeiner, 2015; Cline et al., 2017; Robinson et al., 2020). The finding from our study demonstrates that using multiple sources of information regarding adequate fishing pressure from multiple connected fisheries can also mitigate the effects of resource population fluctuations on harvesting levels. However, our scenarios show that the portfolio effect is only effective when both patches are exhibiting sustainable harvest practices. In the case study, patch 1 was being fished sustainably and patch 2 was experiencing over-fishing, and also included social hierarchy by increasing *ρ*_1_, or the social influence that human population 2 has on the human population in the first patch (table 1). Despite human population 1’s efforts to maintain fish stocks, the unsustainable practices of human population 2 drives the whole fishery to collapse.

We then tested the effect of external social influence (*ρ*) on the case study model and how increasing social influence between human groups would influence the model’s dynamics. Contrary to our previous findings, increasing *ρ* did not result in higher fish populations (Figure 4). Fish populations crashed when *ρ* passed a tipping point, showing that high levels of cooperation between groups resulted in the over-harvest of both populations of fish. At high levels of external social influence, sustainable fishing practices were not achieved because the only information being passed on to the other human population is the number of fishers as opposed to what sustainable fishing practices were used in order to achieve sustainable fishing yields. As a result, when one patch *i* is over-fished and the other patch *j* is fished sustainably, the group *i* will continue to over-fish their own resources because the opposite patch *j* is influencing this group to continue fishing through the high external social influence (*ρ*). Instead of modeling a cohesive system where communication fostered effective conservation, we created a scenario where each community raced to fish each patch as opposed to coming to common understanding of sustainable fishing practices, further highlighting that the content of the information being disseminated matters in successful conservation (Gray et al., 2012). Previous social-ecological research shows that social structures should be taken into consideration when the community manages a resource or else that community management is prone to fail (Grafton, 2005; Newman & Dale, 2007; Cinner et al., 2012; Bodin et al., 2014). Unsuccessful co-management can occur because people who interact differently with the environment or within a society have to consider different trade-offs in conservation, and these trade-offs must be understood in order to institute sustainable practices (Cumming et al., 2017; Baker-Médard, Concannon, et al., 2021). The portfolio effect benefits harvested resources only if each group is participating in sustainable practices.

Further, because of the outside human influence term, *ρ*_*i*_, people are not responding directly to their respective fishing patch, but also to the conservation opinion of the other group. The inclusion of the movement term from each patch overcame the non-linear components of the model because movement is a linear term in this model. Adding a spatial component to socio-ecological models can greatly change their dynamics and therefore how people are expected to act under certain environmental conditions. The dispersion of fish populations must be well understood in order to institute effective conservation practices because any decision made by one group of people to conserve resources may be rendered ineffective if this species is highly migratory and the other group of harvesters is using unsustainable conservation practices. Further, because of the outside influences from the other human patch, fishers are no longer responding directly to fish levels in their respective patch, *i*, but are also influenced by the proportion of fishers in the other patch, *j*. In a scenario where fish is abundant in one patch, this will also encourage fishing in the other patch because incentive to fish will increase from the outside influence parameter. Past research has exemplified how multi-patch models and the addition of spatial components change the dynamics of systems, especially in fisheries (Mchich et al., 2000; Cai et al., 2008; Moeller & Neubert, 2015; Auger et al., 2022).

The decision to include the external social influence term in our model within the injunctive social norms *X*(1 − *X*) implies that external influence can still change an opinion for or against conservation. However, an individual’s willingness to take up a new opinion is still dictated by the overall opinion of their own population exemplifies homophily. Homophily is a concept from sociology where humans tend to take information and the opinions from subgroups similar to them before listening to subgroups of different social standing (Brechwald & Prinstein, 2011). Social network based conservation, like in our model, can replace ‘topdown’ regulation which can exclude stakeholders but has been shown to be susceptible to homophily (Newman & Dale, 2007). Conservation has been shown to be more effective when human populations are more cohesive and that those with subgroups experience more barriers to effective conservation (Bodin & Crona, 2009). Solutions to a lack of cohesion could be to institute some form of liaison that serves as cross-group communicators (Guerrero et al., 2015).

Further research on the model used in this study could consider an open system, where fish diffusion does not necessarily have to pass between patches and could diffuse into non-fished areas. Further, extensions of this work could observe model dynamics with fish species with a long lifespan or fast reproduction rates. Also, stronger social ties have been shown to be more adaptable to environmental change (Grafton, 2005), therefore further studies could evaluate the effect of climate change or extreme events on this social system (White & Wulfing, 2023). The specific way we chose to incorporate social hierarchy into the model could be changed. There are many ways to model social systems so another application of this study would be to compare its results to models that incorporate social hierarchy differently. Next, further work on parameterizing our model to a real-world system could help understand if our model is properly capturing the underlying dynamics of two-patch fishing systems with social hierarchy. Our model only incorporates public opinion, fishing rates, and financial gains from fisheries as aspects that could cause fishery failure. In practice, other issues such as non-compliance to fishing regulations, hyper-stability, and regulation lag time could all be additional factors that result in fishery collapse but are not incorporated in this model (Erisman et al., 2011; Pinsky & Fogarty, 2012; Belhabib et al., 2014). Further, this study does not consider Allee effects in the fish populations, which may alter how spatial dynamics interacts with management practices (White et al., 2021). Finally, our model assumed that the uptake of opinions happens solely through social networks and weighing costs of conservation against the benefits. In reality, there may be more factors that influence one’s harvesting decisions such as governing bodies or media consumption.

## Acknowledgements

This research was supported in part by NSF grant #1923707.

