## Supplemental Material for "Social-ecological models with social hierarchy and spatial structure applied to small scale fisheries"

Equations 2), 4), and 5) are as follows:

$$\frac{dX_i}{dt} = k_i X_i (1 - X_i)[U_{A,i} - U_{B,i}] \tag{2}$$

$$U_{A,i} = \frac{1}{(F_i + c_i)} + d_i X_i + \rho_i X_j \tag{4}$$

$$U_{B,i} = \omega_i + d_i(1 - X_i) + \rho_i(1 - X_j) \tag{5}$$

Substituting equations 4) and 5) into equation 2 Gives:

$\frac{dX_i}{dt} = k_i X_i (1 - X_i)[\frac{1}{(F_i+c_i)} + d_i X_i + \rho_i X_j - \omega_i - d_i(1 - X_i) - \rho_i(1 - X_j)]$

$\frac{dX_i}{dt} = k_i X_i (1 - X_i)[\frac{1}{(F_i+c_i)} - \omega_i + d_i(X_i - 1 + X_i) + \rho_i(X_j - 1 + X_j)]$

$\frac{dX_i}{dt} = k_i X_i (1 - X_i)[\frac{1}{F_i+c_i} - \omega_i + d_i(2X_i - 1) + \rho_i(2X_j - 1)]$
